# Mice lacking triglyceride synthesis enzymes in adipose tissue are resistant to diet-induced obesity

**DOI:** 10.1101/2022.05.05.490833

**Authors:** Chandramohan Chitraju, Alexander W. Fischer, Yohannes A. Ambaw, Kun Wang, Bo Yuan, Sheng Hui, Tobias C. Walther, Robert V. Farese

**Affiliations:** Department of Molecular Metabolism, Harvard T.H. Chan School of Public Health, Boston, MA 02115, USA; Department of Cell Biology, Harvard Medical School, Boston, MA 02115, USA; Broad Institute of Harvard and MIT, Cambridge, MA 02142, USA; Howard Hughes Medical Institute, Boston, MA 02115, USA; Cell Biology Program, Sloan Kettering Institute, Memorial Sloan Kettering Cancer Center, New York, NY, USA

**Author notes:** Correspondence should be addressed to: Tobias C. Walther and Robert V. Farese, Jr. and. These authors contributed equally (Tobias C. Walther, Robert V. Farese).

## Abstract

Triglycerides (TG) in adipocytes provide the major stores of metabolic energy in the body. Optimal amounts of TG stores are desirable as insufficient capacity to store TG, as in lipodystrophy, or exceeding the capacity for storage, as in obesity, results in metabolic disease. We hypothesized that mice lacking TG storage in adipocytes would result in excess TG storage in cell types other than adipocytes and severe lipotoxicity accompanied by metabolic disease. To test this hypothesis, we selectively deleted both TG-synthesis enzymes, DGAT1 and DGAT2, in adipocytes (ADGAT DKO mice). As expected with depleted energy stores, ADGAT DKO mice did not tolerate fasting well and, with prolonged fasting, entered torpor. However, ADGAT DKO mice were unexpectedly otherwise metabolically healthy and did not accumulate TGs ectopically or develop associated metabolic perturbations, even when fed a high-fat diet. The favorable metabolic phenotype resulted from activation of energy expenditure, in part via BAT activation and beiging of white adipose tissue. Thus, the ADGAT DKO mice provide a fascinating new model to study the coupling of metabolic energy storage to energy expenditure.

## INTRODUCTION

Because energy sources are not always available from the environment, many metazoan organisms have evolved the ability to store large amounts of metabolic energy as triglycerides (TG) in adipose tissue. TG is particularly optimal for energy storage because it serves as stores of highly reduced carbon and does not require water for its storage. In cells, such as adipocytes, TGs are stored in organelles called lipid droplets (LDs). In adipocytes of mammals, TGs are stored in a unilocular adipocyte that fills the majority of the aqueous cytoplasm. Although TGs can also be found in LDs in other cell types (i.e., myocytes, hepatocytes, enterocytes), adipocytes represent by far the major energy depots in mammals.

Abundant evidence from many studies suggests that there is an optimal range for adipocyte TG storage in an organism. Exceeding the capacity to store TG in adipocytes occurs in obesity and is often accompanied by deposition of TG in other tissues and metabolic diseases, such as diabetes mellitus or non-alcoholic fatty liver disease. Conversely, insufficient TG storage such as occurs in lipodystrophy is usually associated with adipocyte endocrine deficiency and similar metabolic derangements.

Here, we sought to test the requirement for TG storage in adipocytes in murine physiology at baseline and in response to high-fat feeding. We generated mice lacking both known TG synthesis enzymes, DGAT1 and DGAT2 ^1, 2^, in adipocytes. We expected to generate a mouse model similar to those of classic lipodystrophy due to defects of TG storage in adipose tissue. Moreover, we hypothesized that these mice would have accumulation of TGs in other tissues, such as the liver or skeletal muscle, resulting in lipotoxicity and metabolic derangements, such as insulin resistance or fatty liver disease. To our surprise, we found the opposite result. We report here that selectively impairing TG storage in adipocytes leads to a unique murine model in which depletion of energy stores is not accompanied by metabolic derangements but instead results in protection from adverse metabolic effects, even with high-fat diet (HFD) feeding, due to activation of energy dissipation pathways.

## RESULTS

### ADGAT DKO mice have reduced fat mass and triglycerides in adipose tissue

To generate mice lacking TGs in adipose tissue (ADGAT DKO), we crossed adipose tissue-specific *Dgat1* knockout mice (Cre-transgene expressed under control of the mouse adiponectin promoter ^3^) with *Dgat2* flox mice ^4^. Validation of gene knockouts showed mRNA levels of *Dgat1* were decreased by ∼95% and *Dgat2* by ∼90% in both inguinal white adipose tissue (iWAT) and interscapular brown adipose tissue (BAT) (***Figure 1–figure supplement 1A***). Western blot analysis showed that DGAT1 and DGAT2 proteins were absent in iWAT and BAT (***Figure 1–figure supplement 1B***). *In vitro* DGAT activity in lysates of adipose tissue of ADGAT DKO mice was decreased by ∼80% in iWAT and ∼95% in BAT (***Figure 1–figure supplement 1C***). Similarly, *in vitro* DGAT activity in isolated adipocytes of iWAT was decreased by ∼80% (***Figure 1–figure supplement 1D***).

ADGAT DKO mice appeared healthy (***Figure 1A***) and yielded offspring with the predicted Mendelian ratio of genotypes. Nuclear magnetic resonance imaging showed that fat depots were decreased in chow-fed ADGAT DKO mice (***Figure 1B***). Body weights of 12-week-old chow-fed control mice (*Dgat1* and *Dgat2* double-flexed mice, D1D2 flox) and ADGAT DKO mice were similar (***Figure 1C***), but dual-energy X-ray absorptiometry (DEXA) analysis revealed that the fat mass was decreased by ∼60% in ADGAT DKO mice. The reduction in fat mass was persistent: DEXA analysis of 1-year-old ADGAT DKO mice showed that the fat mass was reduced by ∼75% and that lean mass was increased by ∼15% (***Figure 1–figure supplement 1E***). Visceral white adipose tissue depots (gonadal, mesenteric, pericardial, and perirenal fat depots) were markedly atrophied in ADGAT DKO mice (***Figure 1D***, ***Figure 1–figure supplement 1F***). Gonadal adipose tissue (gWAT) and subcutaneous inguinal WAT (iWAT) in ADGAT DKO mice appeared distinctly “beige” in color (***Figure 1D***, ***Figure 1– figure supplement 1F***). In ADGAT DKO mice, iWAT and BAT were denser, as demonstrated by their sinking in a liquid fixative (***Figure 1F***).

**Figure 1.**
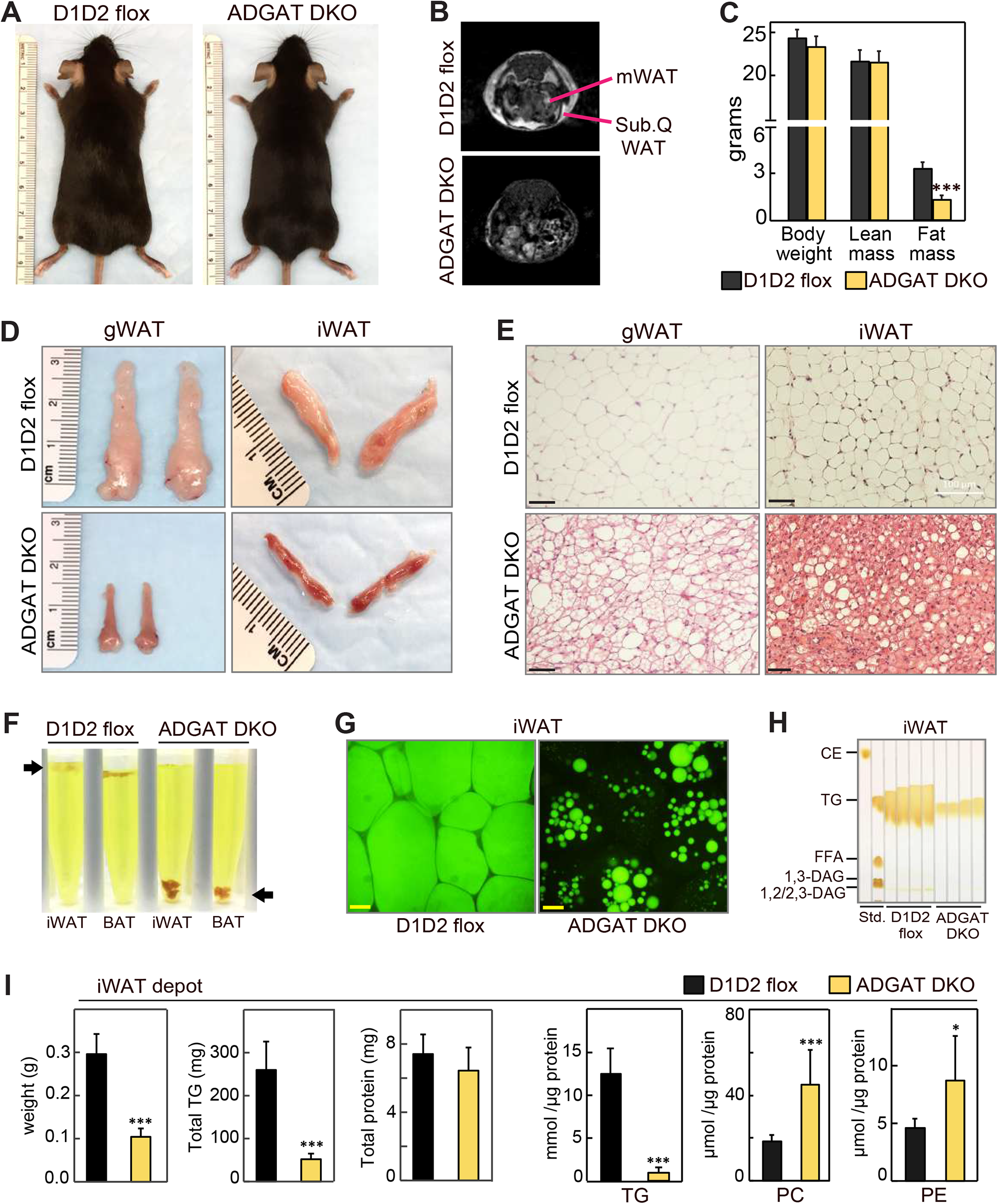
ADGAT DKO mice have reduced fat mass and triglycerides in adipose tissue. (A) Chow-diet-fed ADGAT DKO mice appear normal. Representative photographs of control and ADGAT DKO female mice fed a chow diet. (B) Fat depots were decreased in ADGAT DKO mice. Nuclear magnetic resonance imaging of chow-diet-fed male mice. (C) ADGAT DKO mice have decreased fat mass. Dual-energy X-ray absorptiometry (DEXA) analysis of lean mass and fat mass of chow-diet-fed male mice (n=8). (D) Fat depots were atrophied in ADGAT DKO mice. Representative photographs of iWAT and gWAT from male mice (n=8). (E) gWAT and iWAT of ADGAT DKO mice contain multilocular lipid droplets in adipocytes. H&E-stained sections of gWAT and iWAT from chow-diet-fed male mice (n=6). Scale bars, 50 µm. (F) WAT and BAT of ADGAT DKO mice were denser than controls and sink in an aqueous buffer with fixative (1.25% formaldehyde, 2.5% glutaraldehyde and 0.03% picric acid in 0.1 M sodium cacodylate buffer, pH 7.4, density = 1.01 g/mL) used to fix tissue for electron microscopy. (G) LDs in iWAT of ADGAT DKO mice stain with BODIPY. Confocal fluorescence microscopy images of adipose tissue. LDs were stained by BODIPY 493/503. Scale bar, 25 µm. (H) WAT of ADGAT DKO mice contain triglycerides. Thin layer chromatography analysis of lipids from iWAT of male mice (n=4). TG, triglycerides; CE, cholesterol esters, FFA, free fatty acids; DAG, diacylglycerol. (I) Increased phospholipid levels in iWAT of ADGAT DKO mice. Lipids in iWAT were extracted and quantified by mass spectrometry (n=8). Data are presented as mean ± SD. *p<0.05, ***p<0.001.

The interscapular BAT depot in ADGAT DKO mice appeared darker brown than that in control mice (***Figure 1–figure supplement 2A***). TGs and lipid droplets (LD) were undetectable in BAT of ADGAT DKO mice (***Figure 1–figure supplement 2B,C***). Positron emission tomography-computed tomography scanning using ^18^-fluoro-deoxyglucose (^18^-FDG-PET/CT) showed that, after injection of β-3-adrenoceptor agonist (CL 316,243), BAT of chow-fed ADGAT DKO mice took up more glucose than BAT of control mice (***Figure 1–figure supplement 2D***), presumably to fuel thermogenesis. In agreement with increased glucose uptake, glycogen levels in BAT of ADGAT DKO mice were increased in all conditions except 4°C cold exposure (***Figure 1–figure supplement 2E***). The latter condition may reflect increased glycogen requirements in BAT of ADGAT DKO mice to maintain thermogenesis. This phenotype of DGAT-deficient BAT exhibiting increased glucose uptake and glycogen stores as an alternate fuel is consistent with our previous findings of BAT-specific knockout of both DGAT enzymes ^5^.

### ADGAT DKO mice gradually activate an alternative mechanism to synthesize and store triglycerides

DGAT1 and DGAT2 appear to account for most of TG synthesis in mice. Newborn mice lacking both DGAT enzymes have >95% reduction in whole body TGs ^6^, and adipocytes derived from fibroblasts lacking both enzymes fail to accumulate TGs or LDs ^7^. In agreement with this, histological analysis of WAT of 8-week-old ADGAT DKO mice showed fewer and much smaller LDs than control WAT (***Figure 1E***). However, by age 15 weeks, ADGAT DKO mice exhibited more LDs in iWAT than 8-week-old ADGAT DKO mice, suggesting that they activate alternate pathways to accumulate neutral lipids (***Figure 1–figure supplement 3A***). This finding was more prominent in iWAT than in BAT. The neutral lipid that accumulated was BODIPY-positive (***Figure 1G***), and TLC analysis of adipose tissue lipids from 15-week-old ADGAT DKO mice revealed the lipids to be TGs (***Figure 1H***). Feeding ADGAT DKO mice a HFD increased levels of TGs by ∼twofold in iWAT at age 15 weeks, but the TG content of iWAT remained ∼70% less than control mice (***Figure 1–figure supplement 3A, Figure 4–figure supplement 1A***). The mass reduction of iWAT fat pads was accounted for predominantly by a decrease in TG mass per fat pad (***Figure 1I***); protein levels per fat pad were similar to controls, and no other neutral lipids were detected. Lipid analyses of iWAT by mass spectrometry revealed that TG levels were reduced by ∼80% across all detected TG species (***Figure 1I***, ***Figure 1–figure supplement 3C***). In contrast, several phospholipids were substantially increased in iWAT of ADGAT DKO mice (***Figure 1I***), which may have contributed to the residual fat mass of the iWAT fat pads. Enzyme assays revealed that adipose tissues and isolated white adipocytes from 15-week-old chow-diet-fed ADGAT DKO mice had detectable (∼20% of normal) DGAT activity in iWAT that was not inhibited by DGAT1-or DGAT2-specific inhibitors (***Figure 1–figure supplement 1C,D***). These data suggest that deletion of DGAT1 and DGAT2 in WAT induces a DGAT activity from an alternative enzyme, possibly from other candidates in the DGAT2 gene family ^8^. mRNA levels for MGAT1 and MGAT2, enzymes in the same protein family as DGAT2, were increased in iWAT of ADGAT DKO mice and thus these proteins are candidates for this activity (***Figure 1–figure supplement F, G***).

### Adipose tissue TG stores are required to maintain activity and body temperature during fasting

Because ADGAT DKO mice have severely decreased TG stores, we expected that they would not tolerate fasting well. After 14 hours of fasting, 15-week-old ADGAT DKO mice had lost 10% of body weight (vs. control mice) (***Figure 2A***) and entered a torpor-like state with decreased physical activity and huddling together (***Figure 2–figure supplement movie 1***). Fasting of ADGAT DKO mice also resulted in hypothermia, with body temperatures dropping to ∼30°C (***Figure 2B***), a phenotype exacerbated by cold exposure (***Figure 2E,F***). This differs from the phenotype of BAT DGAT DKO mice, which maintain body temperature during fasting ^5^, presumably because energy stores in WAT are present. Fasting levels of ketone bodies were ∼10% lower, and glucose levels were moderately higher in ADGAT DKO mice than control mice (***Figure 2C,D***), possibly reflecting their greater dependency on glucose as fuel. Thus, as expected, deletion of TG stores in adipose tissue resulted in reduced fuel stores that dramatically altered the physiological responses of the mice to fasting or cold.

**Figure 2.**
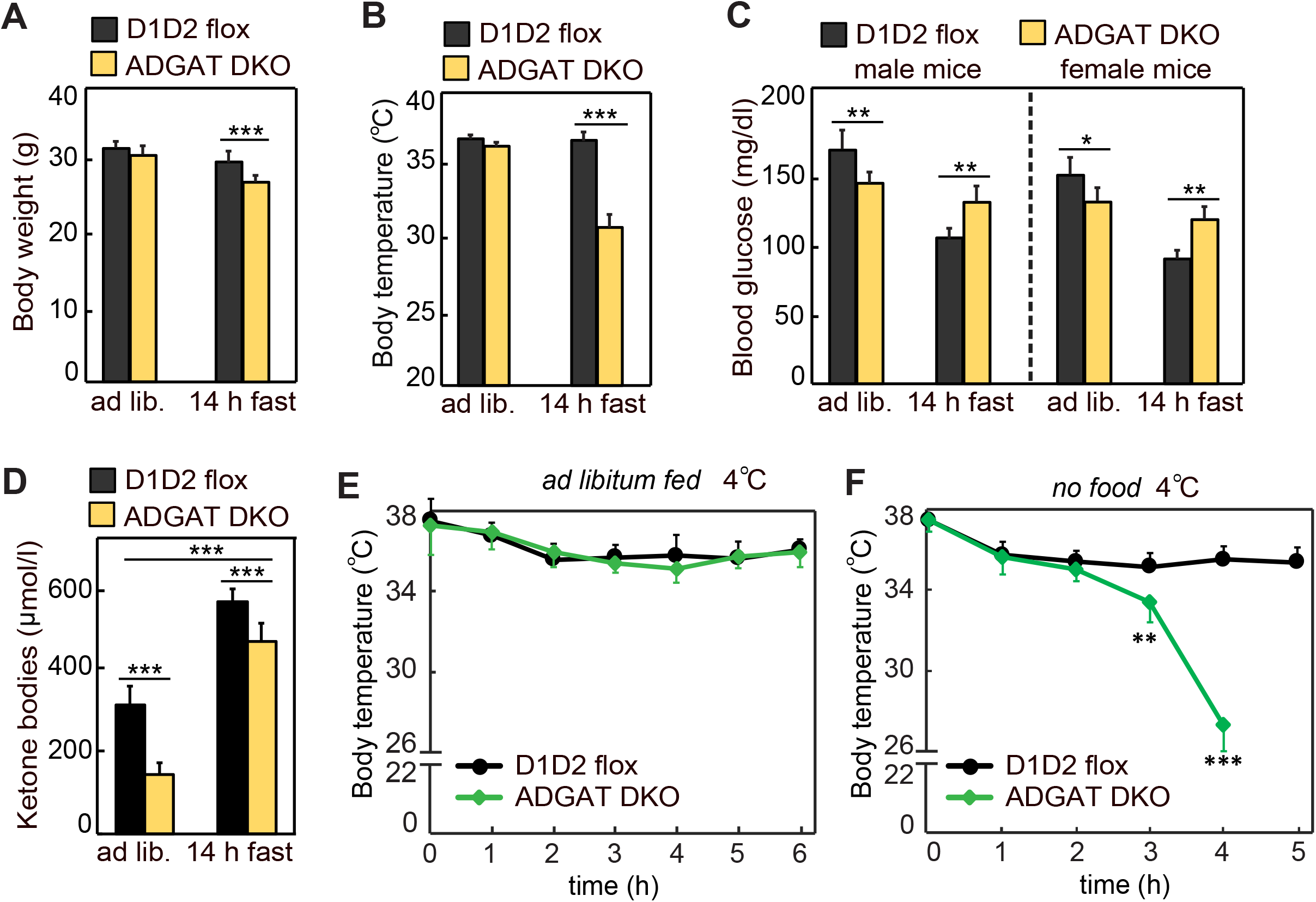
Adipose tissue TG stores are required to prevent a torpor-like state during fasting. (A) Body weights of male mice fed *ad libitum* or fasted 14 h (n=10). Ad lib., *ad libitum* fed. (B) Core body temperature of male mice housed at room temperature and fed *ad libitum* or fasted for 14 h (n=10). (C) Blood glucose levels in male and female mice fed *ad libitum* or fasted for 14 h (n=8). (D) Levels of plasma ketone bodies in male mice fed *ad libitum* or fasted for 14 h (n=8). (E and F) Core body temperature of male mice housed in cold (5° Celsius) with or without food (n=10). Data are presented as mean ± SD. *p<0.05, **p<0.01, ***p<0.001.

### Endocrine function of WAT is maintained in ADGAT DKO mice

Lipodystrophy is a metabolic disease characterized by altered fat distribution, often with severely reduced amounts of adipose tissue and TG storage. Classically, lipodystrophy in humans and mice is accompanied by reduced levels of adipocyte-derived endocrine hormones and often results in insulin resistance and diabetes ^9–11^. ADGAT DKO mice share an impaired capacity to store TG in adipose tissue with lipodystrophy models. However, despite this, in these mice adiponectin and leptin mRNA levels were moderately increased in iWAT (***Figure 3A***), whereas the mRNA levels of *Plin1* were unchanged (***Figure 3A***), and plasma levels of adiponectin and leptin were normal and 40% decreased, respectively (***Figure 3B***). When normalized to adipose tissue weight, leptin levels were similar to control mice (***Figure 3C***). Glucose levels in ADGAT DKO mice fasted for 4 h were slightly lower (169 ± 16 mg/dl vs. 144 ± 14 mg/dl, respectively, p<0.01) than in control mice, and insulin levels were not different (***Figure 3D***). Analysis of plasma metabolites showed a ∼30% reduction in non-esterified fatty acids, a ∼15% reduction in glycerol (***Figure 3E***), and a ∼50% reduction in ketones in chow-diet-fed ADGAT DKO mice (***Figure 3F***). Glucose and insulin tolerances were similar in chow-diet-fed ADGAT DKO and control mice (***Figure 3G,H***). Thus, despite the impairment of fat storage in adipose tissue, ADGAT DKO mice had substantial levels of adipocyte-derived endocrine hormones and apparently normal glucose metabolism.

**Figure 3.**
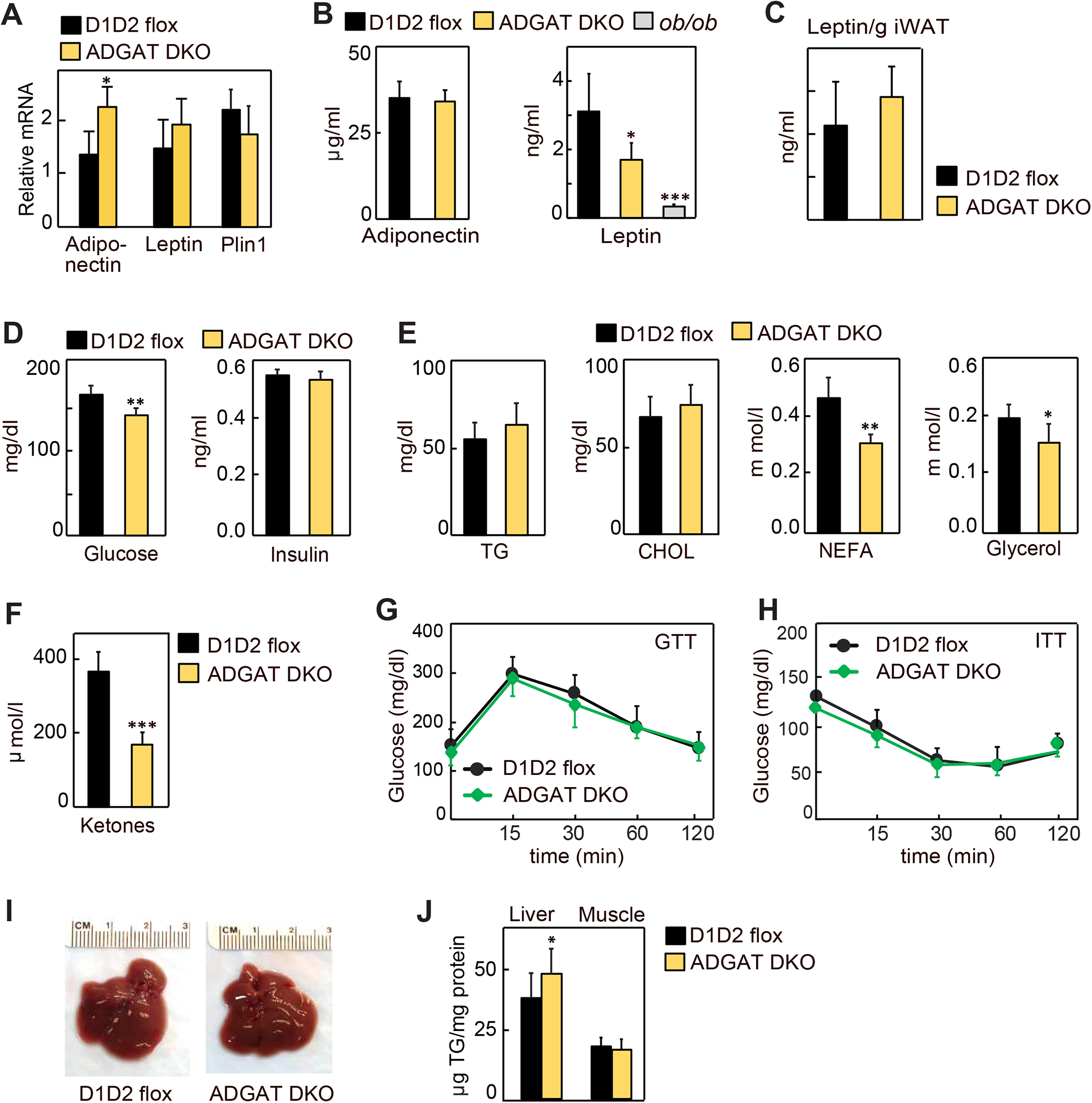
Lipodystrophy is uncoupled from detrimental metabolic effects in ADGAT DKO mice. (A) Adiponectin and leptin mRNA levels were moderately increased in iWAT of ADGAT DKO mice. Relative mRNA levels of leptin and adiponectin in iWAT of chow-diet-fed male mice (n=6). (B) Plasma levels of adiponectin were normal, and leptin levels were moderately decreased in *ad libitum* chow-diet-fed male mice (n=8). (C). Plasma leptin levels normalized per gram of WAT mass (n = 8). (D) ADGAT DKO mice had normal glucose and insulin levels. Glucose and insulin levels in *ad libitum* chow-diet-fed male mice (n=8). (E) Decreased free fatty acids in ADGAT DKO mice. Levels of plasma metabolites in *ad libitum* chow-diet-fed male mice (n=8). (F) Decreased ketones in *ad libitum* chow-diet-fed ADGAT DKO male mice (n=8). (G and H) Glucose- and insulin-tolerance tests were normal in chow-diet-fed male mice (n=10). (I) ADGAT DKO mice had non-steatotic livers. Representative photographs of livers from chow-diet-fed mice. (J) Triglyceride were moderately increased in livers of ADGAT DKO mice. Triglyceride levels in livers and skeletal muscle of chow-diet-fed male mice (n=6). Data are presented as mean ± SD. *p<0.05, **p<0.01, ***p<0.001.

Lipodystrophy is also typically accompanied by ectopic lipid deposition, particularly manifesting as hepatic steatosis ^12–14^. In contrast, livers of ADGAT DKO mice appeared normal (***Figure 3I***), with moderately increased weights in 15-week-old chow-diet-fed mice (1.7 ± 0.2 g vs. 2.1 ± 0.3 g, p<0.05). TG levels were only modestly increased (∼10%) in livers of ADGAT DKO mice and were unchanged in skeletal muscle (***Figure 3J***). Thus, ADGAT DKO mice, with markedly reduced TG storage in adipocytes, were remarkably metabolically healthy, with essentially none of the metabolic derangements typically associated with lipodystrophy.

### ADGAT DKO mice are resistant to diet-induced obesity and associated metabolic derangements

We next tested whether the metabolically heathy phenotype of ADGAT DKO mice would persist with feeding of a western-type high-fat diet (HFD), which normally causes obesity and insulin resistance. We hypothesized that fatty acids from the HFD would not be stored in adipocytes of ADGAT DKO mice and as a consequence ectopically accumulate, resulting in tissue lipotoxicity. However, after feeding ADGAT DKO mice a HFD for 12 weeks, they appeared healthy and remained relatively lean, with both male and female mice gaining ∼40% less body weight than control D1D2 flox mice (***Figure 4A–C***). The reduction in body weight was due to a ∼70% reduction in fat mass (***Figure 4C***). Food intake during HFD feeding was similar (***Figure 4D***), implying ADGAT DKO mice have increased energy expenditure. This was validated by indirect calorimetry, where ADGAT DKO mice exhibited increased energy expenditure that was particularly prominent during night-time, when the mice were eating (***Figure 4E***). The respiratory exchange ratio (RER) was lower in ADGAT DKO mice during HFD feeding (***Figure 4F***), consistent with increased fat oxidation. We did not measure caloric loss in the feces of ADGAT DKO mice and would not expect this with adipocyte-specific deletions of DGAT1 and DGAT2.

**Figure 4.**
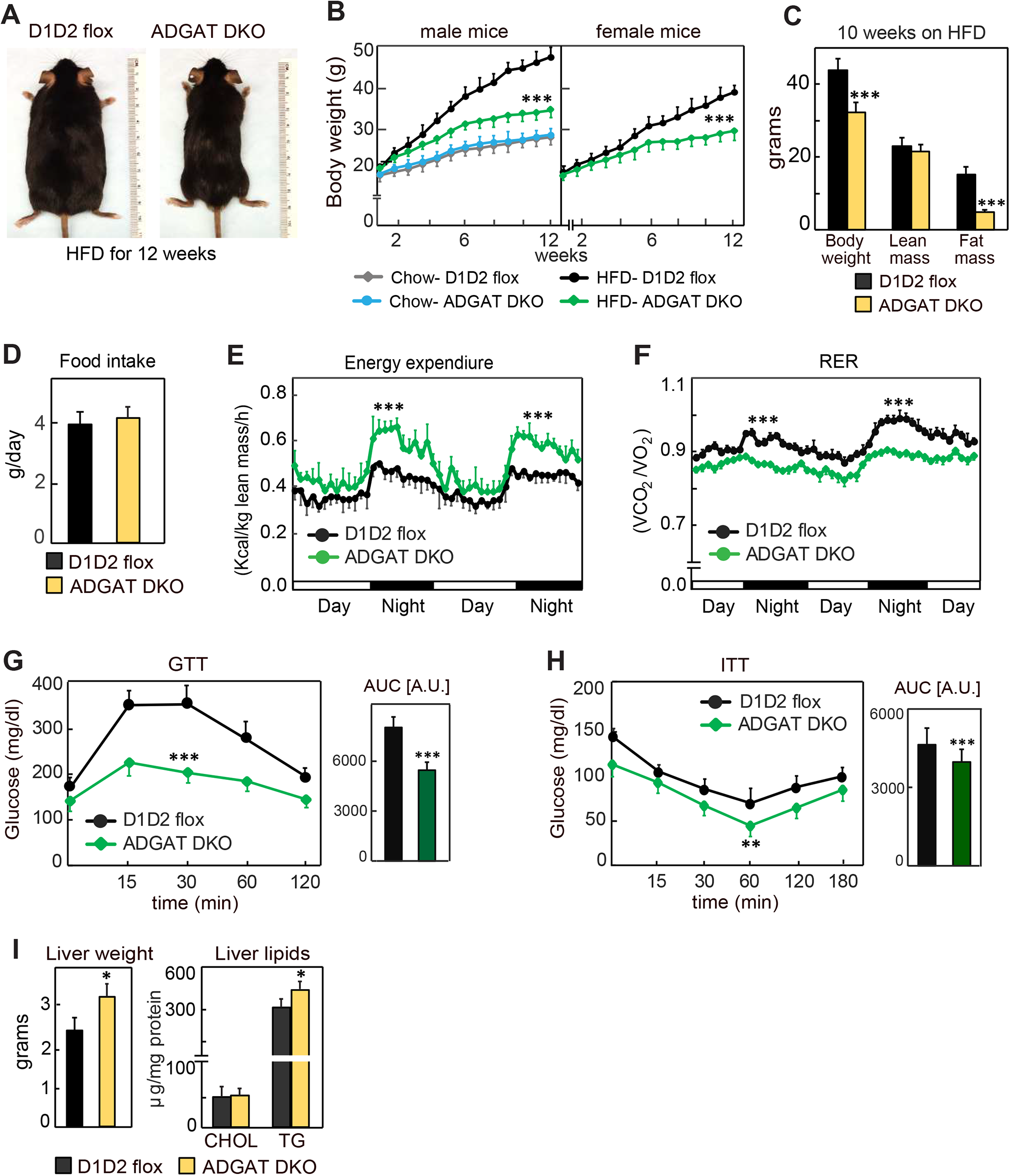
ADGAT DKO mice are resistant to diet-induced obesity and glucose intolerance. (A) ADGAT DKO mice stay lean on an HFD. Representative photographs of male mice fed on HFD for 12 weeks. (B) Both male and female ADGAT DKO mice gained ∼40% less body weight than control mice. Body weights of mice fed on a chow-diet or HFD (n=15 for males, n=12 for females). (C) ADGAT DKO mice had decreased fat mass on HFD feeding. DEXA analysis of lean mass and fat mass of HFD fed male mice (n=10). (D) ADGAT DKO male mice had normal food intake during HFD feeding (n=5). (E and F) ADGAT DKO mice had increased energy expenditures. Energy expenditure and respiratory quotient on HFD-fed male mice measured by indirect calorimetry. Values were normalized to lean mass (n=4). (G and H) ADGAT DKO mice were protected from HFD-induced glucose intolerance and insulin resistance. Glucose- and insulin-tolerance tests were performed on HFD-fed (for 9 or 10 weeks, respectively) male mice (n=10). AUC; area under the curve. (I) Liver weights and triglyceride levels were moderately increased in HFD-fed ADGAT DKO male mice (n=6). Data are presented as mean ± SD. *p<0.05, **p<0.01, ***p<0.001.

We also examined metabolic parameters in the HFD-fed ADGAT DKO mice. Plasma glucose levels were slightly lower in ADGAT DKO, and insulin levels were not different (***Figure 4–figure supplement 1B,C***). ADGAT DKO mice were protected from HFD-induced glucose intolerance (***Figure 4G***). The insulin response was similar in ADGAT DKO and control mice, although the ADGAT DKO mice basal levels of glucose were reduced (***Figure 4H***). Liver weights and hepatic TG levels were markedly increased with HFD in both ADGAT DKO mice and controls and were ∼20% and ∼10% higher in ADGAT DKO mice, respectively (***Figure 4I***). Hepatic cholesterol levels were similar (***Figure 4I***). HFD-induced activation of ER-stress response in the livers was similar to control mice (***Figure 4–figure supplement 1D***). Thus, surprisingly, despite not being able to robustly store TGs in adipocytes, ADGAT DKO mice were resistant to most effects of an HFD, and our studies indicate that they activate compensatory mechanisms of energy expenditure that increase fat oxidation.

### ADGAT DKO mice activate energy dissipation mechanisms, including adipocyte beiging in WAT

We next investigated the mechanisms for improved metabolic health in ADGAT DKO mice. Browning or beiging of adipose tissue is associated with improved metabolic health in mice and humans ^15–18^, and the “beige” appearance in iWAT and gWAT depots of ADGAT DKO mice (***Figure 1D,E***) suggested that beiging adaptations may be present. Histological examination showed almost all adipocytes in both iWAT and gWAT contained multi-locular LDs (***Figure 5A,C***). mRNA levels of signature genes of adipocyte beiging, such as *Ucp1* (∼600-fold), *Idea* (∼20-fold), *Pparα* (∼10-fold), and *Pgc1α* (∼sixfold), were markedly increased in iWAT and gWAT of room temperature housed chow-fed ADGAT DKO mice (***Figure 5B,D***). Fatty acid levels were decreased; intermediates of glycolysis and Krebs cycle were enriched in both iWAT and BAT of ADGAT DKO mice, consistent with increased glycolysis and fatty acid oxidation (***Figure 5–figure supplement 1A,B***). Protein levels of UCP1 and respiratory complex proteins were also markedly greater in iWAT of room-temperature-housed chow-fed ADGAT DKO mice than controls and were even further increased under HFD conditions (***Figure 5E***). Beiging of adipocytes in iWAT appeared independent of ambient temperature and was also present in ADGAT DKO mice after 6 weeks of thermoneutral housing (***Figure 5F***), and blood glucose levels were moderately lower in thermoneutral housed male and female mice (***Figure 5G***). Beiging appeared to be non-cell-autonomous, as the changes found in beige fat were largely absent in cells differentiated from pre-adipocytes, with the exception of a twofold increase of *Ucp1* mRNA levels (***Figure 5–figure supplement 2A–D***), suggesting that beiging in ADGAT DKO mice is activated in part through the sympathetic nervous system (SNS) ^19, 20^. Hormones, such as FGF21, also can activate beiging, either via the SNS ^21, 22^ or in a paracrine manner ^21, 23^. FGF21 mRNA levels were increased by ∼twofold and ∼sixfold in liver and iWAT of ADGAT DKO mice, respectively (***Figure 5–figure supplement 3A***), and plasma levels of FGF21 were increased ∼threefold in ADGAT KO mice (***Figure 5–figure supplement 3B***). However, plasma FGF21 levels were similar in ADGAT DKO mice and controls that were fed an HFD, suggesting an endocrine FGF21 effect is not responsible for the increased beiging of iWAT (***Figure 5–figure supplement 3B–D***).

**Figure 5.**
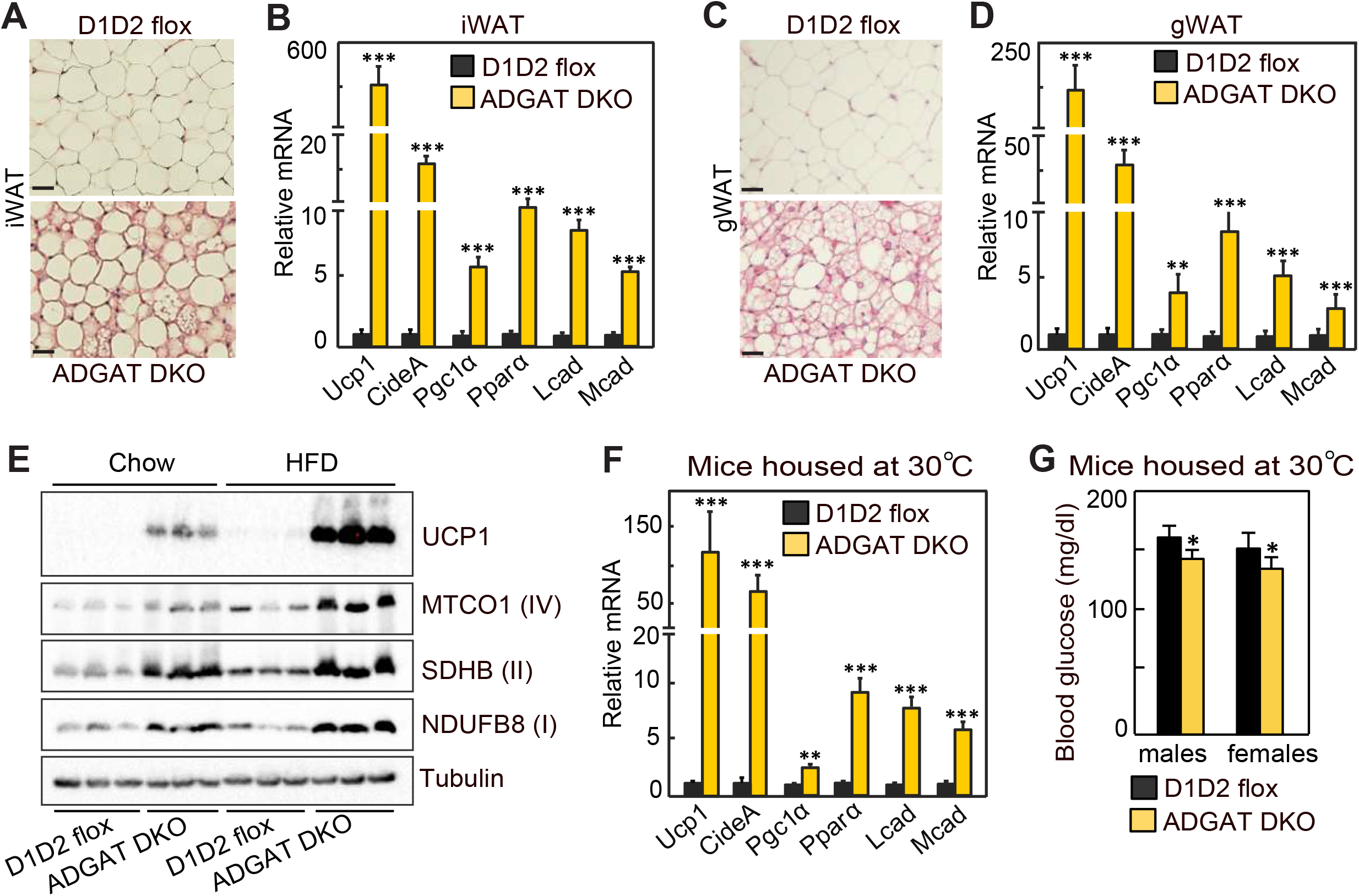
Beiging of WAT in ADGAT DKO mice. (A) ADGAT DKO mice contain multi-locular LDs in adipocytes of iWAT. H&E-stained sections of iWAT from male mice fed a chow diet and housed at room temperature (n=6). Scale bars, 50 µm. (B) Increased expression of thermogenic marker genes in iWAT of ADGAT DKO mice. Relative mRNA levels of thermogenic genes in iWAT of male mice fed a chow diet and housed at room temperature (n=6). (C) ADGAT DKO mice contain multi-locular LDs in adipocytes of gWAT. H&E-stained sections of gWAT from male mice fed a chow diet and housed at room temperature (n=6). Scale bars, 50 µm. (D) Increased expression of thermogenic marker genes in gWAT of ADGAT DKO mice. Relative mRNA levels of thermogenic genes in gWAT of male mice fed a chow diet and housed at room temperature (n=6). (E) HFD feeding increases levels of UCP1 in iWAT of ADGAT DKO mice. Immunoblot analysis of UCP1 and OXPHOS proteins in iWAT of male mice fed either a chow diet or an HFD (n=3). Mice were housed at room temperature. (F) Beiging was intact in iWAT of thermoneutral-housed ADGAT DKO mice. Relative mRNA levels of thermogenic genes in iWAT of chow-diet-fed male mice housed at thermoneutral temperature for 6 weeks (n=6). (G) Blood glucose levels were normal in thermoneutral housed ADGAT DKO mice. Glucose levels in male mice fed a chow diet and housed at thermoneutral temperature for 6 weeks (n=8). Data are presented as mean ± SD. *p<0.05, **p<0.01, ***p<0.001.

## DISCUSSION

We report a novel mouse model with impaired TG synthesis in adipocytes. The resultant defect in TG stores had both expected and surprising effects on murine physiology. First, as expected, ADGAT DKO mice did not tolerate fasting well. At room temperature, fasted ADGAT DKO mice entered a torpor-like state, characterized by decreased ambulation and a drop in body temperature. Torpor is a physiological state that enables conservation of metabolic energy and the signals to induce this state are poorly understood ^24^. The phenotype of ADGAT DKO mice suggests that depletion of adipocyte TG stores is sufficient to induce energy conservation and is somehow sensed.

A surprising feature of this mouse model is the apparent metabolic health, despite the reduced capacity to store lipids in the adipose tissue. Typically, loss of white adipose tissue (WAT) leads to a condition known as lipodystrophy ^25, 26^. Lipodystrophy patients nearly always present with many metabolic derangements, including ectopic lipid accumulation in the liver (hepatic steatosis), hypertriglyceridemia, and insulin resistance or diabetes ^27–29^. A characteristic feature of lipodystrophy is the decrease in levels of adipose tissue-derived hormones, such as leptin ^30, 31^, and many derangements of lipodystrophies are corrected by leptin therapy ^9, 32^.

Remarkably, the lipodystrophic ADGAT DKO mice with marked reductions in fat storage were metabolically healthy and did not develop metabolic derangements, such as diabetes or hepatic steatosis, even when fed an HFD. The metabolic health of the ADGAT DKO mice likely is due to their intact ability to make adipose tissue and to maintain adipose tissue endocrine function. These findings are consistent with previous data showing that cultured adipocytes differentiate normally in the absence of TG storage ^7^. Adipose tissue of ADGAT DKO mice maintained the ability to synthesize and secrete adipose-derived hormones, such as leptin, which is crucially absent in typical lipodystrophy ^32–35^. Leptin levels often correlate with adiposity and TG stores ^36^, but ADGAT DKO mice exhibit a dissociation of TG stores with leptin expression, thereby showing that these parameters appear not to be causally related. Instead, leptin levels may be better correlated with other adipose properties, such as the number of adipocytes, as reflected in the observed correlation of leptin with adipose tissue mass.

Our studies revealed that, in response to the compromised ability to store TG in white and brown adipose tissue, mice activate pathways of energy expenditure, including the generation of beige adipocytes and activation of BAT. Aspects of this phenotype were present even at thermoneutral conditions. The mechanisms underlying the activation of energy expenditure and beiging in ADGAT-DKO mice are presently unclear, but may involve SNS stimulation. The lack of fatty acids in adipocytes may result in other available fuels (fatty acids and glucose) being routed to the BAT and beige adipocytes to maintain body temperature. One possible mechanism for the beiging and increased energy expenditure is that the ADGAT DKO mice secrete a factor (or factors) that activates the SNS and beiging pathway. Although currently such a factor remains to be identified, this model is reminiscent of global DGAT1 knockout mice ^37^, which exhibit increased energy expenditure, enhanced glucose metabolism, protection from diet-induced obesity, and increased leptin sensitivity ^38–40^. For global DGAT1 knockout mice, fat transplant studies suggested the adipose tissue is the source of such factors ^39^.

Hormones that activate beiging, such as FGF21, are also testable candidate factors that may contribute to the ADGAT DKO phenotype. Many of the beneficial metabolic effects that we found in ADGAT DKO mice, including increased energy expenditure, increased glucose uptake by BAT, and torpor and browning, are found in murine models with increased levels of FGF21 ^21, 22, 41, 42^. However, an endocrine effect of FGF21 seems unlikely in these mice due to similar levels of the hormone in plasma of HFD-fed mice. We note, however, that a paracrine effect is not excluded. Crossing ADGAT DKO mice with FGF21 knockout mice would further test FGF21’s contribution to the metabolic phenotype of these mice. It would also be interesting to inhibit both DGAT enzymes at early or late time points in adipocyte differentiation and determine if endocrine or paracrine factors were altered, which might explain the systemic effect on metabolism.

In summary, ADGAT DKO mice represent an intriguing model in which a marked reduction in the ability to store TG in adipocytes triggers organismal pathways of energy dissipation. This suggests that exceeding the capacity to store energy in adipocytes is somehow sensed and triggers thermogenesis in adipose tissue. This phenotype likely requires an intact adipocyte endocrine system, which was found in ADGAT DKO mice but if often deficient in other models of lipodystrophy. The exact mechanism for how a TG storage defect triggers energy dissipation is currently unclear, but unraveling of this mechanism could lead to new strategies for treating or reducing obesity.

## ACKNOWLEDGMENTS

We thank members of the Farese & Walther laboratory for helpful comments and G. Howard for editorial assistance. We thank Karen Inouye and Sarah Mitchell for helping with indirect calorimetry analysis. Nathan Heinzman for helping with metabolomics experiment and the Longwood small animal imaging facility at Beth Israel Deaconess Medical Center for PET/CT analysis. This work was supported in part by NIH grant R01GM124348 (to R.F.). T.C.W is an investigator of the Howard Hughes Medical Institute.

## CONTRIBUTIONS

C.C., R.V.F., and T.C.W. planned the study and designed the experiments. C.C. generated DGAT2 flox, DGAT double flox and ADGAT DKO mice. C.C. performed most of the experiments. A.W.F. performed denervation of mice. K.W. analyzed metabolomics data. Y.A. performed lipidomics analysis. B.Y. and S.H. performed metabolomics. C.C., R.V.F., and T.C.W. wrote the manuscript. All authors read and edited the manuscript.

## STAR★METHODS

- KEY RESOURSES TABLE
- CONTACT FOR REAGENT AND RESOURSE SHARING
- EXPERIMENTAL MODEL AND SUBJECT DETAILS

- Generation of ADGAT DKO mice
- Animal Husbandry
- Cold exposure Studies
- METHOD DETAILS

- DGAT Activity Assay
- Tissue Lipid Analysis
- Microscopy and Image Processing
- [^18^F]-FDG-PET/CT Analysis
- RNA extraction and Quantitative Real-Time PCR
- Immunoblotting
- Comprehensive Lab Animal Monitoring System (CLAMS)
- Denervation of Adipose Tissue
- QUANTIFICAION AND STATISTICAL ANALYSIS

### CONTACT FOR REAGENT AND RESOURCES SHARING

Further information and request for reagents and resources should be mailed to Robert V. Farese, Jr. and Tobias C. Walther.

### EXPERIMENTAL PROCEDURES

#### Generation of ADGAT DKO Mice

To generate adipose tissue–specific *Dgat1* and *Dgat2* double-knockout (ADGAT DKO) mice, we first generated *Dgat1* and *Dgat2* double-floxed mice (D1D2 flox) by crossing *Dgat1^flox/flox^* mice ^43^ (Jackson Laboratory stock number: 017322) with *Dgat2^flox/flox^* mice ^4^ (Jackson Laboratory stock number: 033518). To generate ADGAT DKO mice, we crossed D1D2 flox mice with transgenic mice expressing Cre recombinase under control of the murine adiponectin promoter ^3^ (Jackson Laboratory stock number: 028020).

#### Animal Husbandry

All mouse experiments were performed under the guidelines from Harvard Center for Comparative Medicine. Mice were maintained in a barrier facility, at room temperatures (22–23°C), on a regular 12-h light and 12-h dark cycle and had *ad libitum* access to food and water unless otherwise stated. For thermoneutral studies, mice were housed at 29°C. Mice were fed on standard laboratory chow diet (PicoLab^®^ Rodent Diet 20, 5053; less than 4.5% crude fat) or Western-type high-fat diet (Envigo, TD.88137; 21.2% fat by weight, 42% kcal from fat).

#### Cold-Exposure Studies

For cold-exposure experiments (at 5°C), mice were single-housed in the morning around 8:00 am. Mice had free access to food and water unless otherwise stated. Core body temperatures were recorded using a rectal probe thermometer.

#### DGAT Activity Assay

DGAT enzymatic activity was measured in WAT and BAT lysates at V_max_ substrate concentrations. Assay mixture contained 20 µg of adipose tissue lysate, 100 µM of 1,2-dioleoyl-*sn*-glycerol, 25 µM of oleoyl-CoA, which contained [^14^C] oleoyl-CoA as tracer, and 5 mM MgCl_2_ in an assay in buffer containing 100 mM Tris-HCl (pH 7.4) and protease inhibitors. Reactions were carried out as described ^2, 44^. After stopping the reaction, lipids were extracted and separated by TLC using a hexane:diethyl ether:acetic acid (80:20:1) solvent system. The TLC plates were exposed to phosphor imager screen and developed.

#### Tissue Lipid Analysis

Approximately 50 mg of adipose tissue was homogenized in 1 mL of lysis buffer (250 mM sucrose, 50 mM Tris Cl, pH 7.0, with protease inhibitor cocktail (11873580001, Roche)). The homogenate was mixed with 5 ml of chloroform: methanol (3:2 v:v) and extracted for 2 h by vigorous shaking. Upon centrifugation at 3000 x g at room temperature for 10 min, 100 µL of lower organic phase was collected and dried in a speed vac. To the dried lipids, 100–300 µL of 0.1% Triton X-100 was added, and the solution was sonicated using ultrasonic homogenizer (Biologics, Inc., model 3000MP) for 10 sec with 30% amplitude. The total TG content was measured using the Infinity TM triglycerides reagent (Thermo Scientific) according to the manufacturer’s protocol. TG and total cholesterol were measured using Infinity TM triglycerides reagent (Thermo Scientific) and a cholesterol E kit (Wako Diagnostics), respectively, according to manufacturer’s protocol. For plasma lipids measurement, 5 µL of plasma was used directly.

#### Microscopy and Image Processing

Microscopy was performed on spinning disk confocal microscope (Yokogawa CSU-X1) set up on a Nikon Eclipse Ti inverted microscope with a 100× ApoTIRF 1.4 NA objective (Nikon) in line with 2x amplification. BODIPY 493/503 fluorophore was exited on 561-nm laser line. Fluorescence was detected by an iXon Ultra 897 EMCCD camera (Andor). Acquired images were processed using FIJI software (http://fiji.sc/Fiji).

#### RNA Extraction and Quantitative Real-Time PCR (qRT-PCR)

Total RNA from tissues was isolated with the Qiazol lysis reagent and using the protocol of the RNeasy Kit (Qiagen). Complementary DNA was synthesized using the iScript cDNA Synthesis Kit (Bio-Rad), and qPCRs were performed using the SYBR Green PCR Master Mix Kit (Applied Biosystems).

#### Immunoblotting

Tissues were lysed using RIPA lysis buffer (25 mM Tris Cl, pH 7.6, 150 mM NaCl, 1% NP-40, 1% sodium deoxycholate, 0.1% SDS) containing protease inhibitors (11873580001, Roche). Proteins were denatured in Laemmli buffer and separated on 10% SDS-PAGE gels and transferred to PVDF membranes (Bio-Rad). The membranes were blocked with blocking buffer for 1 h in TBST containing 5% BSA or 5% milk, and then incubated with primary antibodies overnight. The membranes were then washed three times with TBST for 10 min, and incubated in mouse secondary antibodies (Santa Cruz Biotechnology) at 1:5000 dilutions in blocking buffer. Membranes was washed again three times with TBST for 10 min, and revealed using the Super Signal West Pico kit (Thermo Scientific).

#### Comprehensive Lab Animal Monitoring System (CLAMS)

Mice were housed individually and acclimatized for 2 days. Oxygen consumption, carbon dioxide release, energy expenditure, and activity were measured using a Columbus Instruments’ Oxymax Comprehensive Lab Animal Monitoring System (CLAMS) system according to guidelines for measuring energy metabolism in mice ^45^.

#### Lipidomics Analysis

For lipidomic analysis of iWAT, ∼50 mg of iWAT was homogenized in 1 mL ice-cold phosphate-buffered saline using a bead mill homogenizer. Tissue lysates (50 μg) were transferred to a pyrex glass tubes with a PTFE-liner cap. Lipids were extracted by Folch method ^46^, briefly, 6 mL of ice-cold chloroform-methanol (2:1 v/v) and 1.5 mL of water were added to the samples, and tubes were vortexed thoroughly to mix the samples homogenously with a polar and non-polar solvent. SPLAH mix internal standards were spiked in before the extraction. The organic phase of each sample was normalized by total soluble protein amounts and measured by BCA assay (Thermo Scientific, 23225, Waltham, MA). After vortexing, samples were centrifuged for 30 min at 1100 rpm at 4°C to separate the organic and inorganic phases. Using a sterile glass pipette, the lower organic phase was transferred into a new glass tube, taking care to avoid the intermediate layer of cellular debris and precipitated proteins. The samples were dried under nitrogen flow until the solvents were completely dried. Samples were resuspended in 250 μL of chloroform: methanol 2:1 and stored in -80 until mass spectrometer (MS) analysis. Lipids were separated using ultra-high-performance liquid chromatography (UHPLC) coupledv with tandem MS. Briefly, UHPLC analysis was performed on a C30 reverse-phase column (Thermo Acclaim C30, 2.1 x 250 mm, 3 μm operated at 55° C; Thermo Fisher Scientific) connected to a Dionex UltiMate 3000 HPLC system and a QExactive orbitrap MS (Thermo Fisher Scientific) equipped with a heated electrospray ionization probe. 5 μL of each sample was analyzed separately, using positive and negative ionization modes. Mobile phase contained 60:40 water:acetonitrile (v:v), 10 mM ammonium formate and 0.1% formic acid, and mobile phase B consisted of 90:10 2-propanol/acetonitrile, also including 10 mM ammonium formate and 0.1% formic acid. MS spectra of lipids were acquired in full-scan/data-dependent MS2 mode. For the full-scan acquisition, the resolution was set to 70,000, the AGC target was 1e6, the maximum injection time was 50 msec, and the scan range was m/z = 133.4– 2000. For data-dependent MS2, the top 10 ions in each full scan were isolated with a 1.0 Da window, fragmented at a stepped normalized collision energy of 15, 25, and 35 units, and analyzed at a resolution of 17,500 with an AGC target of 2e5 and a maximum injection time of 100 msec. Peak identification and data analysis were carried out using Lipid Search software version 4.1 SP (Thermo Fisher Scientific) ^47^.

#### Metabolomic Analysis

BAT and iWAT was snap frozen in liquid nitrogen and ground at cryogenic temperature with a cyromill (Retsch, Newtown, PA). The tissue was extracted with -20°C 40: 40: 20 methanol: acetonitrile: water at a concentration of 25 mg/mL. Samples were vigorously vortexed and centrifuged at 4 °C at 16,000 g for 10 min, and the supernatant was transferred to LC-MS vials for analysis. Chromatographic separation was performed using XBridge BEH Amide XP Column (2.5 µm, 2.1 mm × 150 mm) with associated guard column (2.5 µm, 2.1 mm X 5 mm) (Waters, Milford, MA). The mobile phase A was 95% water and 5% acetonitrile, containing 10 mM ammonium hydroxide and 10 mM ammonium acetate. The mobile phase B was 80% acetonitrile and 20% water, with 10 mM ammonium hydroxide and 10 mM ammonium acetate. The linear elution gradient was: 0 ∼ 3 min, 100% B; 3.2 ∼ 6.2 min, 90% B; 6.5. ∼ 10.5 min, 80% B; 10.7 ∼ 13.5 min, 70% B; 13.7 ∼ 16 min, 45% B; and 16.5 ∼ 22 min, 100% B. The flow rate was 0.3 mL/ min. The autosampler was maintained at 4°C. The injection volume was 5 µL, and needle wash was performed between samples using 40: 40: 20 methanol: acetonitrile: water. The MS used was Q Exactive HF (Thermo Fisher Scientific, San Jose, CA), and scanned from 70 to 1000 *m/z* with switching polarity. The resolution was 120,000. Metabolites were identified based on accurate mass and retention time using an in-house library, and the software used was EI-Maven (Elucidata, Cambridge, MA). Data was analyzed using R software (version 4.2.0). The ion intensity of each sample was first normalized to the corresponding sample protein content. Differentially abundant metabolites were analyzed with the limma R/Bioconductor package, and the multiple comparisons were corrected using the Benjamini-Hochberg procedure (adjusted *p* value; q value). The volcano plots were generated using the ggplot and ggrepel packages.

#### Statistical Analyses

Data are presented as mean ± SD (standard deviation). Statistical significance was evaluated by unpaired two-tailed Student’s t-test or two-way ANOVA with Bonferroni’s multiple comparison test. Significant differences are annotated as follows: *p < 0.05, **p < 0.01, ***p < 0.001.

## SUPPLEMENTAL INFORMATION

### FIGURE LEGENDS FOR SUPPLEMENTAL FIGURES

**Figure 1—figure supplement 1.**
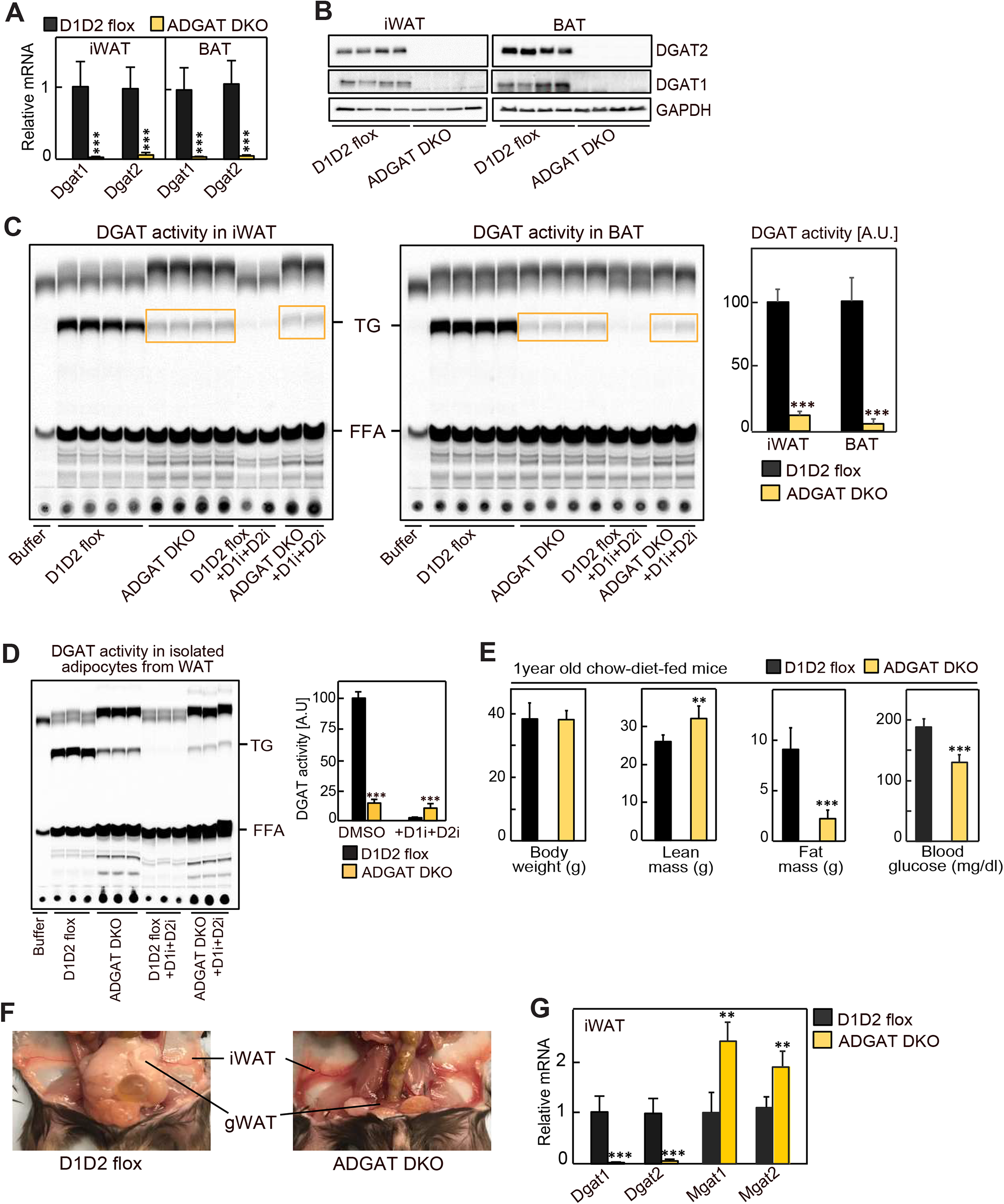
(A) *Dgat1* and *Dgat2* transcripts levels were decreased in iWAT and BAT of ADGAT DKO mice. Relative mRNA levels in iWAT and BAT of male mice fed a chow diet (n=6). (B) DGAT1 and DGAT2 proteins were absent in iWAT and BAT of ADGAT DKO mice. Western blot analysis of DGAT1 and DGAT2 in iWAT and BAT of male mice fed a chow diet (n=4). (C) *In-vitro* DGAT activity was decreased in lysates of iWAT and BAT of ADGAT DKO male mice (n=4). (D) *In-vitro* DGAT activity was decreased in isolated adipocytes from iWAT of ADGAT DKO male mice (n=4). (E) Body weights, lean mass and fat mass analysis of 1-year-old male mice fed a chow diet (n=10). (F) Gross appearance of WAT depots in male mice. **p<0.01, ***p<0.001. (G) Relative mRNA levels of MGAT enzymes in iWAT of male mice fed a chow diet (n=6).

**Figure 1—figure supplement 2.**
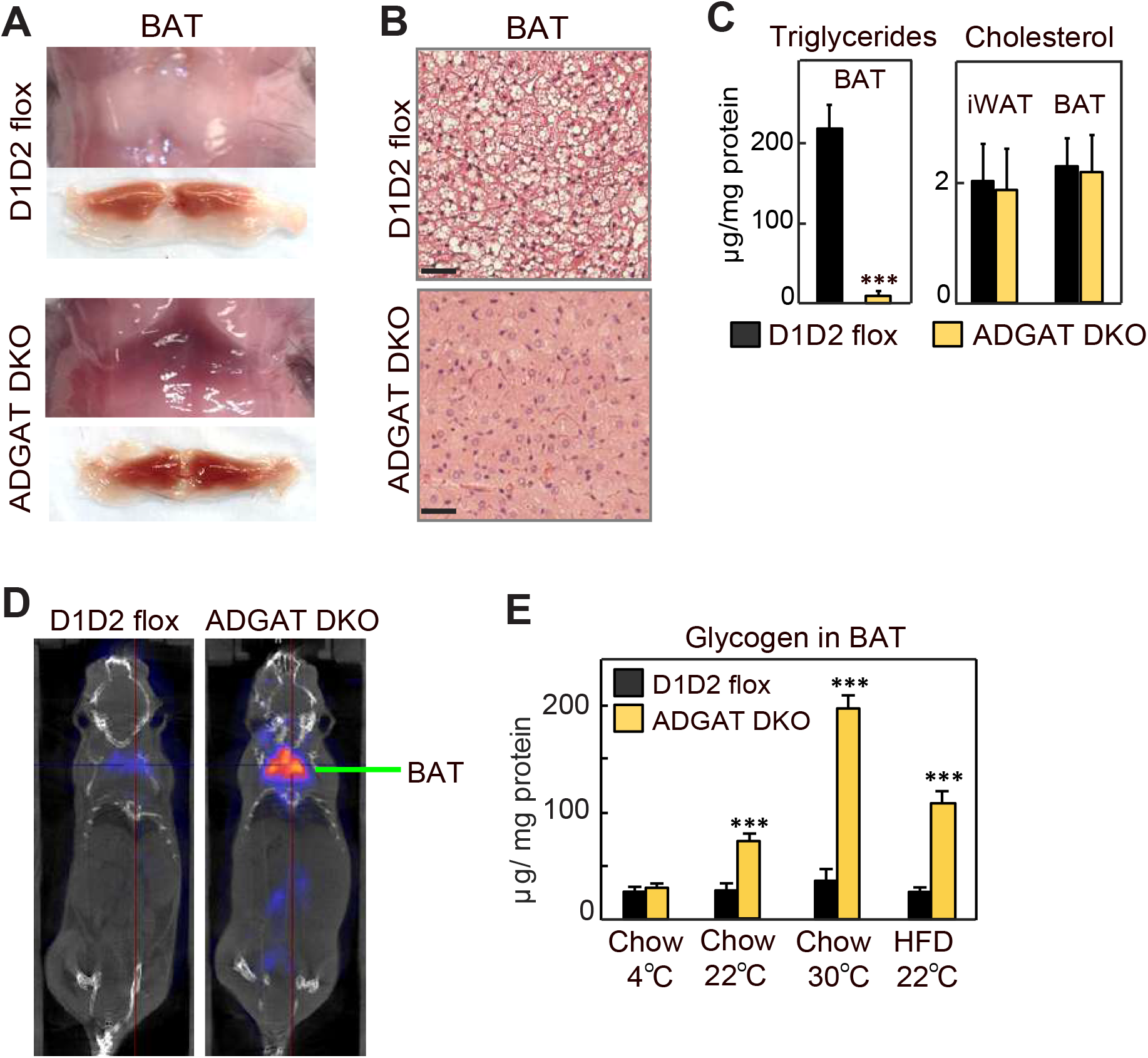
(A) Gross appearance of BAT. (B) H&E-stained sections of iBAT (n=6). Scale bars, 50 µm. (C) Triglycerides and total cholesterol levels in BAT (n=6). (D) [^18^F]-FDG-PET/CT scans of male mice administered CL 316,243. (E) Glycogen levels in iBAT of male mice (n=6). Data are presented as mean ± SD. ***p<0.001.

**Figure 1—figure supplement 3.**
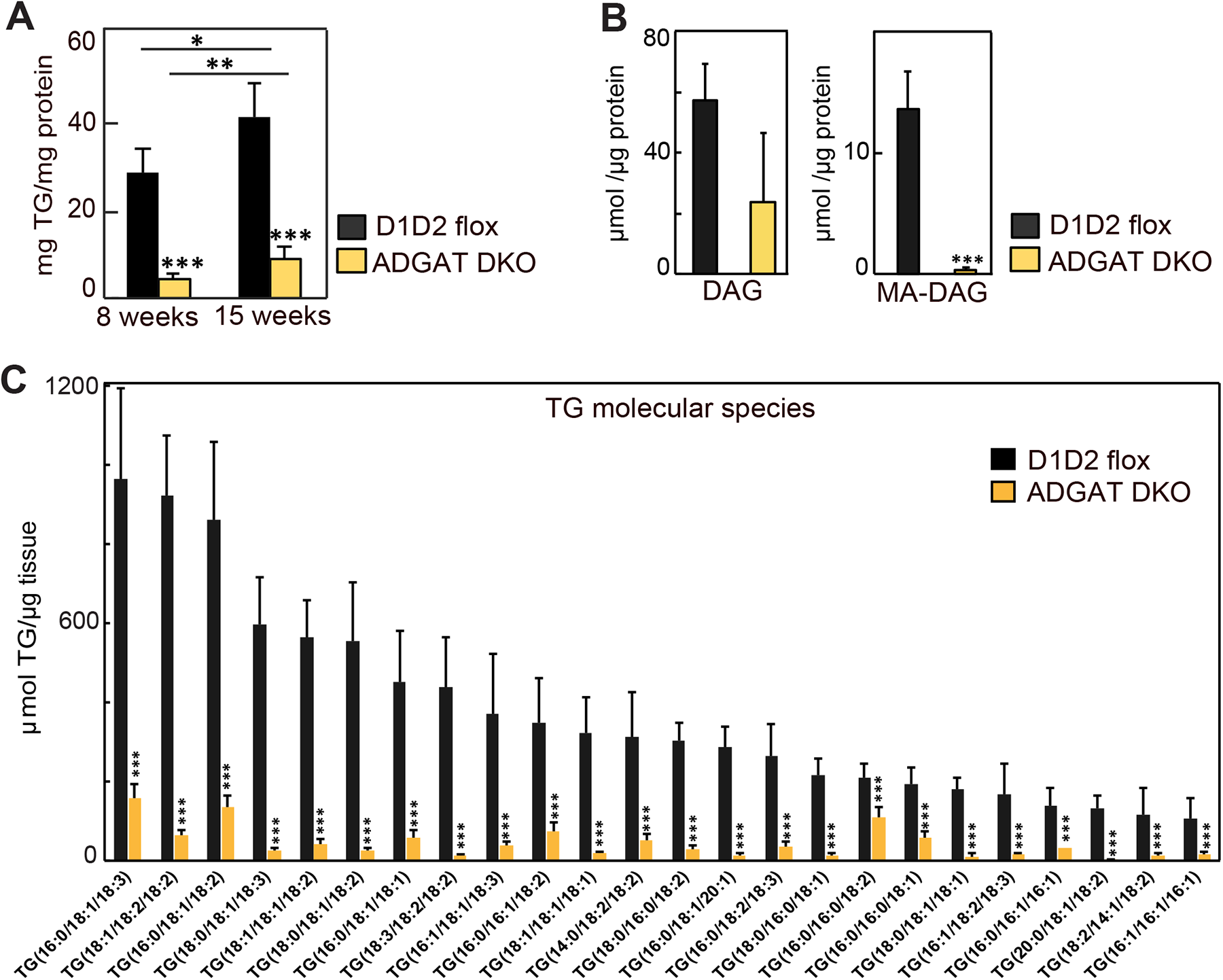
(A) Triglycerides were decreased in iWAT of ADGAT DKO male mice (n=6). Triglyceride levels were measured using Infinity triglyceride reagent. (B) Diacylglycerol and mono alkyl-diacylglycerol were decreased in iWAT of ADGAT DKO mice (n=6). Lipids were quantified by mass spectrometry. (C) Triglyceride molecular species were decreased in iWAT of ADGAT DKO mice (n=6). Lipids were quantified by mass spectrometry. Data are presented as mean ± SD. *p<0.05, **p<0.01, ***p<0.001.

**Figure 4—figure supplement 1.**
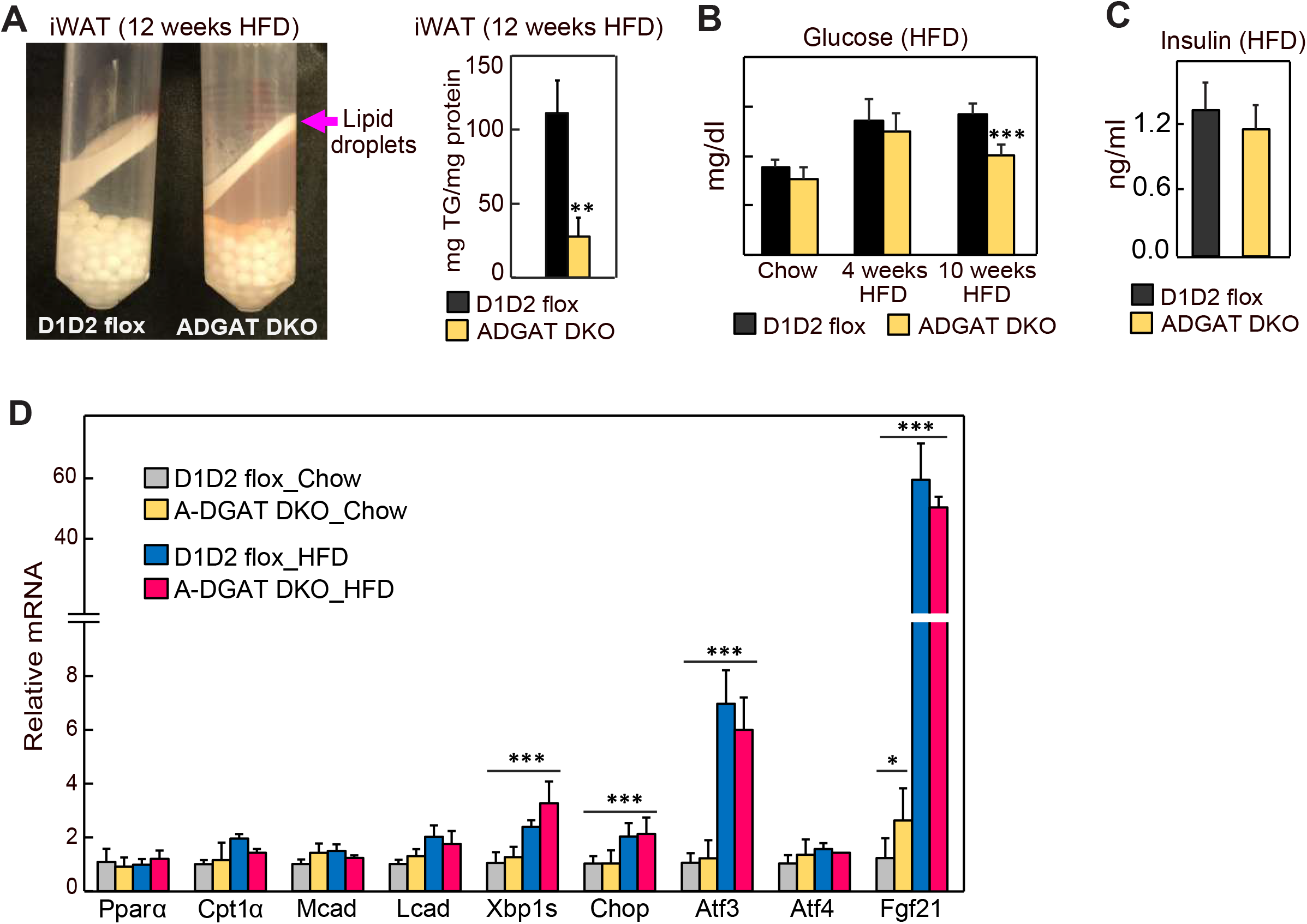
(A) LDs isolated from iWAT of mice fed an HFD. (B) Blood glucose levels in high-fat-diet fed male mice (n=8). (C) Insulin levels in high-fat-diet fed male mice (n=8). (D) Relative mRNA levels in livers of HFD mice (n=6). Data are presented as mean ± SD. *p<0.05, **p<0.01, ***p<0.001.

**Figure 5—figure supplement 1.**
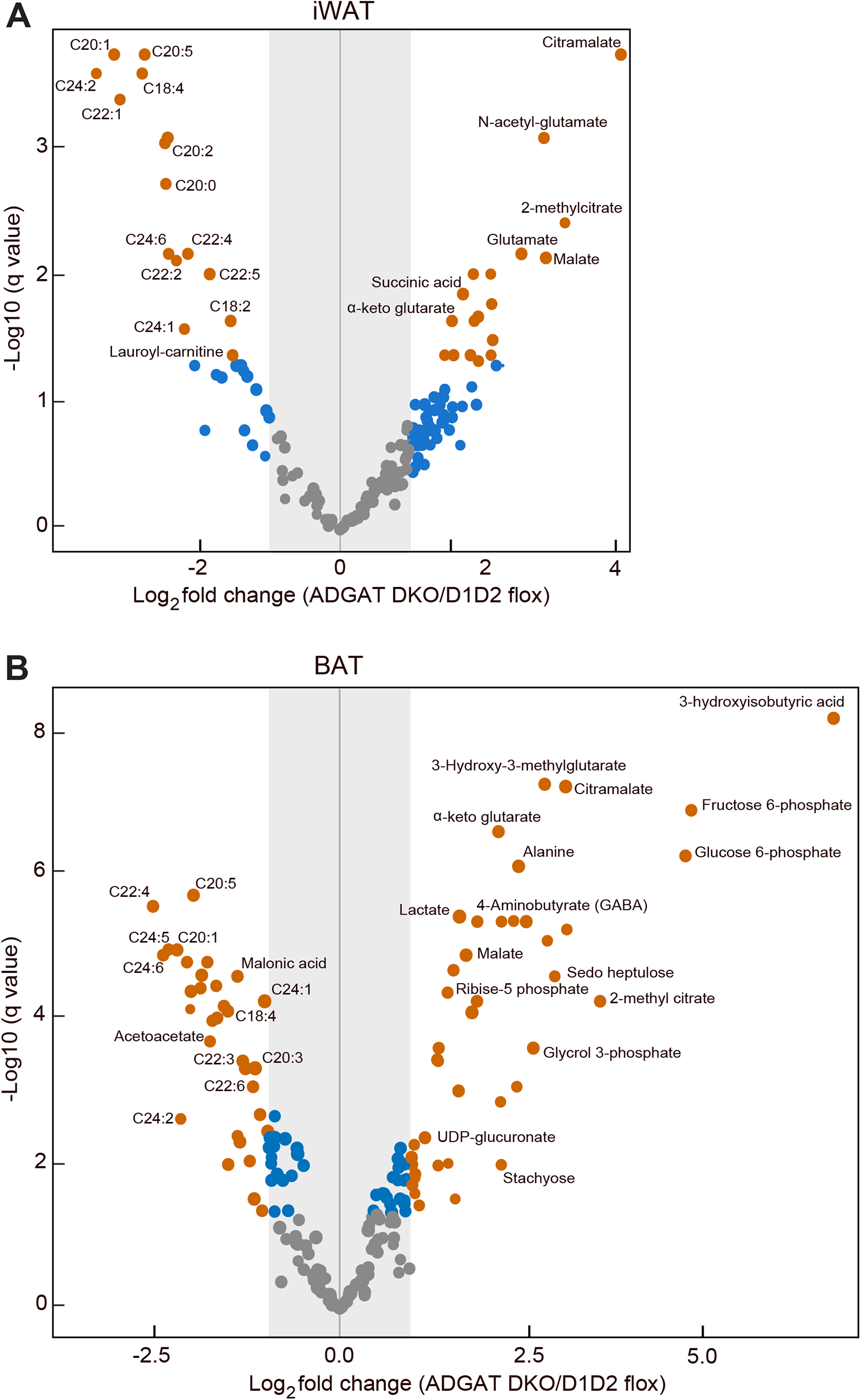
(A) Volcano plot showing differentially abundant metabolites in iWAT of chow-diet-fed male mice (n=8). (B) Volcano plot showing differentially abundant metabolites in iBAT of chow-diet-fed male mice (n=8). Orange dots represent metabolites with more than twofold change (adjusted p values or q<0.05)). Blue dots represent metabolites with more than twofold change but not statistically significant. Grey dots represent metabolites that were unchanged between control and ADGAT DKO mice.

**Figure 5—figure supplement 2.**
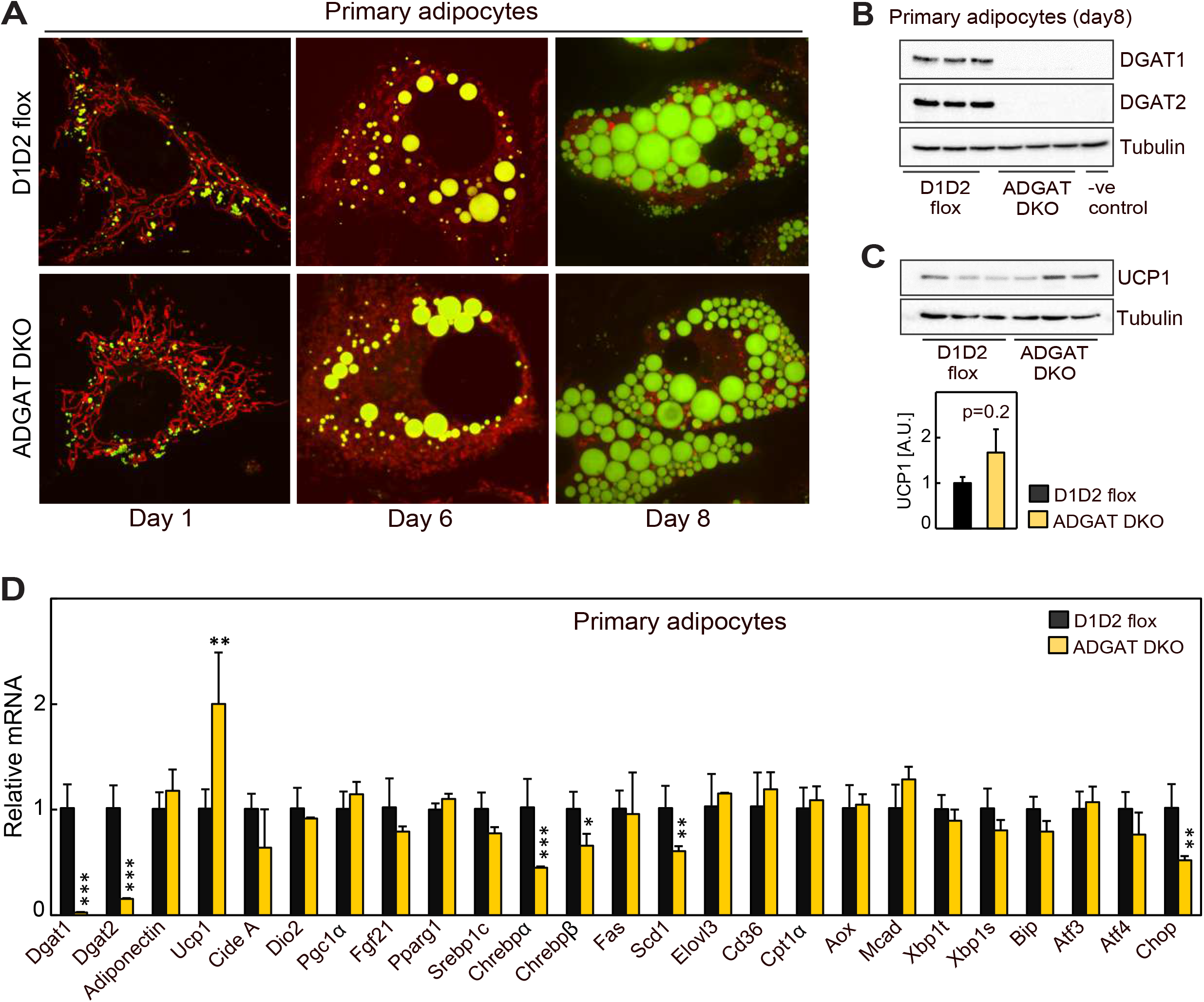
(A) Primary adipocytes differentiated from stromal vascular fraction of iWAT from ADGAT DKO male mice contain lipid droplets. LDs were stained by BODIPY 493/503. (B and C) Western blot analysis of DGATs and UCP1 in primary adipocytes (n=3). (D) Relative mRNA levels in day-8 primary adipocytes loaded with oleic acid for 16 h (n=3). Data are presented as mean ± SD. *p<0.05, **p<0.01, ***p<0.001.

**Figure 5—figure supplement 3.**
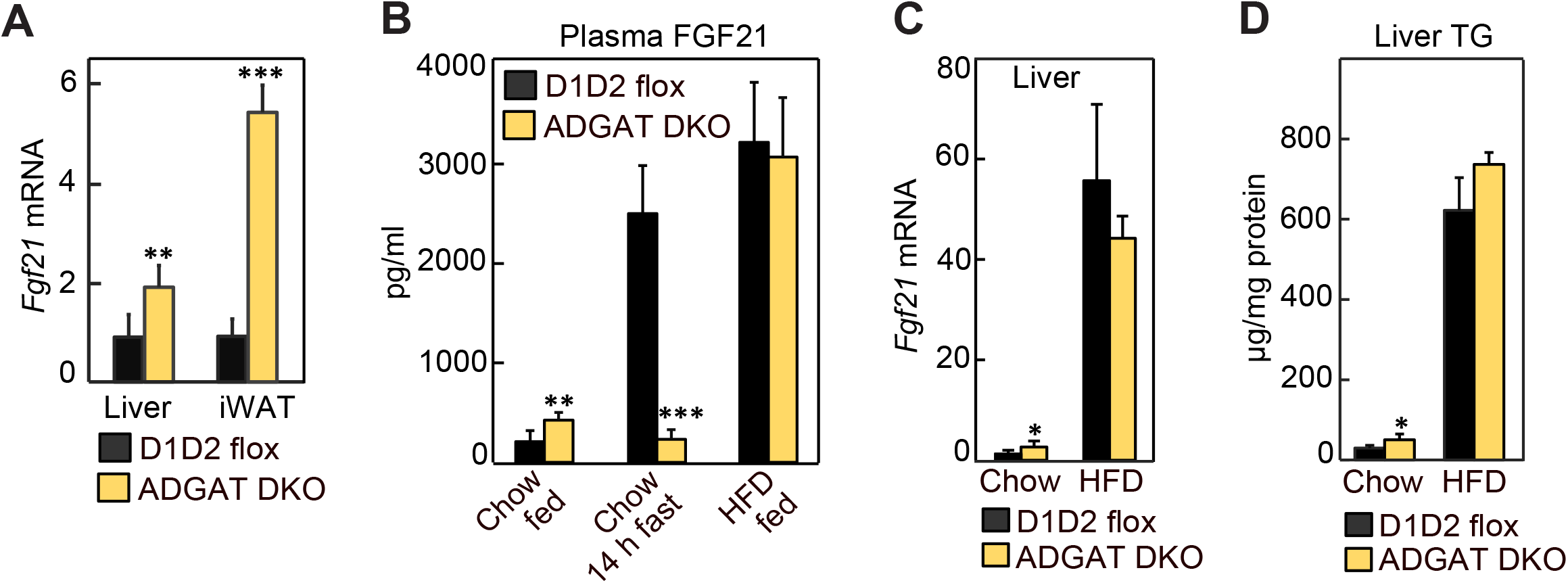
(A) FGF21 transcript levels were increased in iWAT and livers of ADGAT DKO mice. Relative mRNA levels of FGF21 in iWAT and livers of chow-diet fed male mice (n=6). (B) FGF21 levels were increased in ADGAT DKO mice. Plasma levels of FGF21 in chow-diet-fed (*ad libitum* fed or 14 h fasted) or HFD-fed male mice (n=8). (C) FGF21 transcript levels in livers of chow-diet- or HFD-fed male mice (n=6). (D) Triglyceride levels in livers of chow-diet- or HFD-fed male mice (n=6). Data are presented as mean ± SD. *p<0.05, **p<0.01, ***p<0.001.

## REFERENCES

1. Cases, S., Smith, S.J., Zheng, Y.W., Myers, H.M., Lear, S.R., Sande, E., Novak, S., Collins, C., Welch, C.B., Lusis, A.J., et al. (1998). Identification of a gene encoding an acyl CoA:diacylglycerol acyltransferase, a key enzyme in triacylglycerol synthesis. Proc Natl Acad Sci U S A 95, 13018–13023.

2. Cases, S., Stone, S.J., Zhou, P., Yen, E., Tow, B., Lardizabal, K.D., Voelker, T., and Farese, R.V., Jr. (2001). Cloning of DGAT2, a second mammalian diacylglycerol acyltransferase, and related family members. J Biol Chem 276, 38870–38876. 10.1074/jbc.M106219200.

3. Eguchi, J., Wang, X., Yu, S., Kershaw, E.E., Chiu, P.C., Dushay, J., Estall, J.L., Klein, U., Maratos-Flier, E., and Rosen, E.D. (2011). Transcriptional control of adipose lipid handling by IRF4. Cell Metab 13, 249–259. 10.1016/j.cmet.2011.02.005.

4. Chitraju, C., Walther, T.C., and Farese, R.V., Jr. (2019). The triglyceride synthesis enzymes DGAT1 and DGAT2 have distinct and overlapping functions in adipocytes. J Lipid Res 60, 1112–1120. 10.1194/jlr.M093112.

5. Chitraju, C., Fischer, A.W., Farese, R.V., Jr., and Walther, T.C. (2020). Lipid Droplets in Brown Adipose Tissue Are Dispensable for Cold-Induced Thermogenesis. Cell Rep 33, 108348. 10.1016/j.celrep.2020.108348.

6. Stone, S.J., Myers, H.M., Watkins, S.M., Brown, B.E., Feingold, K.R., Elias, P.M., and Farese, R.V., Jr. (2004). Lipopenia and skin barrier abnormalities in DGAT2-deficient mice. J Biol Chem 279, 11767–11776. 10.1074/jbc.M311000200.

7. Harris, C.A., Haas, J.T., Streeper, R.S., Stone, S.J., Kumari, M., Yang, K., Han, X., Brownell, N., Gross, R.W., Zechner, R., and Farese, R.V., Jr. (2011). DGAT enzymes are required for triacylglycerol synthesis and lipid droplets in adipocytes. J Lipid Res 52, 657–667. 10.1194/jlr.M013003.

8. Yen, C.L., Brown, C.H.t., Monetti, M., and Farese, R.V., Jr. (2005). A human skin multifunctional O-acyltransferase that catalyzes the synthesis of acylglycerols, waxes, and retinyl esters. J Lipid Res 46, 2388–2397. 10.1194/jlr.M500168-JLR200.

9. Oral, E.A., Simha, V., Ruiz, E., Andewelt, A., Premkumar, A., Snell, P., Wagner, A.J., DePaoli, A.M., Reitman, M.L., Taylor, S.I., et al. (2002). Leptin-replacement therapy for lipodystrophy. N Engl J Med 346, 570–578. 10.1056/NEJMoa012437.

10. Reue, K., and Phan, J. (2006). Metabolic consequences of lipodystrophy in mouse models. Curr Opin Clin Nutr Metab Care 9, 436–441. 10.1097/01.mco.0000232904.82038.db.

11. Peterfy, M., Phan, J., Xu, P., and Reue, K. (2001). Lipodystrophy in the fld mouse results from mutation of a new gene encoding a nuclear protein, lipin. Nat Genet 27, 121–124. 10.1038/83685.

12. Cui, X., Wang, Y., Tang, Y., Liu, Y., Zhao, L., Deng, J., Xu, G., Peng, X., Ju, S., Liu, G., and Yang, H. (2011). Seipin ablation in mice results in severe generalized lipodystrophy. Hum Mol Genet 20, 3022–3030. 10.1093/hmg/ddr205.

13. Moitra, J., Mason, M.M., Olive, M., Krylov, D., Gavrilova, O., Marcus-Samuels, B., Feigenbaum, L., Lee, E., Aoyama, T., Eckhaus, M., et al. (1998). Life without white fat: a transgenic mouse. Genes Dev 12, 3168–3181. 10.1101/gad.12.20.3168.

14. Shimomura, I., Hammer, R.E., Richardson, J.A., Ikemoto, S., Bashmakov, Y., Goldstein, J.L., and Brown, M.S. (1998). Insulin resistance and diabetes mellitus in transgenic mice expressing nuclear SREBP-1c in adipose tissue: model for congenital generalized lipodystrophy. Genes Dev 12, 3182–3194. 10.1101/gad.12.20.3182.

15. Becher, T., Palanisamy, S., Kramer, D.J., Eljalby, M., Marx, S.J., Wibmer, A.G., Butler, S.D., Jiang, C.S., Vaughan, R., Schoder, H., et al. (2021). Brown adipose tissue is associated with cardiometabolic health. Nat Med 27, 58–65. 10.1038/s41591-020-1126-7.

16. Wu, J., Bostrom, P., Sparks, L.M., Ye, L., Choi, J.H., Giang, A.H., Khandekar, M., Virtanen, K.A., Nuutila, P., Schaart, G., et al. (2012). Beige adipocytes are a distinct type of thermogenic fat cell in mouse and human. Cell 150, 366–376. 10.1016/j.cell.2012.05.016.

17. van Marken Lichtenbelt, W.D., Vanhommerig, J.W., Smulders, N.M., Drossaerts, J.M., Kemerink, G.J., Bouvy, N.D., Schrauwen, P., and Teule, G.J. (2009). Cold-activated brown adipose tissue in healthy men. N Engl J Med 360, 1500–1508. 10.1056/NEJMoa0808718.

18. Kazak, L., Chouchani, E.T., Jedrychowski, M.P., Erickson, B.K., Shinoda, K., Cohen, P., Vetrivelan, R., Lu, G.Z., Laznik-Bogoslavski, D., Hasenfuss, S.C., et al. (2015). A creatine-driven substrate cycle enhances energy expenditure and thermogenesis in beige fat. Cell 163, 643–655. 10.1016/j.cell.2015.09.035.

19. Kajimura, S., Spiegelman, B.M., and Seale, P. (2015). Brown and Beige Fat: Physiological Roles beyond Heat Generation. Cell Metab 22, 546–559. 10.1016/j.cmet.2015.09.007.

20. Harms, M., and Seale, P. (2013). Brown and beige fat: development, function and therapeutic potential. Nat Med 19, 1252–1263. 10.1038/nm.3361.

21. Fisher, F.M., Kleiner, S., Douris, N., Fox, E.C., Mepani, R.J., Verdeguer, F., Wu, J., Kharitonenkov, A., Flier, J.S., Maratos-Flier, E., and Spiegelman, B.M. (2012). FGF21 regulates PGC-1alpha and browning of white adipose tissues in adaptive thermogenesis. Genes Dev 26, 271–281. 10.1101/gad.177857.111.

22. Owen, B.M., Ding, X., Morgan, D.A., Coate, K.C., Bookout, A.L., Rahmouni, K., Kliewer, S.A., and Mangelsdorf, D.J. (2014). FGF21 acts centrally to induce sympathetic nerve activity, energy expenditure, and weight loss. Cell Metab 20, 670–677. 10.1016/j.cmet.2014.07.012.

23. Abu-Odeh, M., Zhang, Y., Reilly, S.M., Ebadat, N., Keinan, O., Valentine, J.M., Hafezi-Bakhtiari, M., Ashayer, H., Mamoun, L., Zhou, X., et al. (2021). FGF21 promotes thermogenic gene expression as an autocrine factor in adipocytes. Cell Rep 35, 109331. 10.1016/j.celrep.2021.109331.

24. Gavrilova, O., Leon, L.R., Marcus-Samuels, B., Mason, M.M., Castle, A.L., Refetoff, S., Vinson, C., and Reitman, M.L. (1999). Torpor in mice is induced by both leptin-dependent and -independent mechanisms. Proc Natl Acad Sci U S A 96, 14623–14628. 10.1073/pnas.96.25.14623.

25. Mann, J.P., and Savage, D.B. (2019). What lipodystrophies teach us about the metabolic syndrome. J Clin Invest 129, 4009–4021. 10.1172/JCI129190.

26. Patni, N., and Garg, A. (2015). Congenital generalized lipodystrophies--new insights into metabolic dysfunction. Nat Rev Endocrinol 11, 522–534. 10.1038/nrendo.2015.123.

27. Lawrence, R.D. (1946). Lipodystrophy and hepatomegaly with diabetes, lipaemia, and other metabolic disturbances; a case throwing new light on the action of insulin. (concluded). Lancet 1, 773. 10.1016/s0140-6736(46)91599-1.

28. Misra, A., and Garg, A. (2003). Clinical features and metabolic derangements in acquired generalized lipodystrophy: case reports and review of the literature. Medicine (Baltimore) 82, 129–146. 10.1097/00005792-200303000-00007.

29. Reitman, M.L. (2002). Metabolic lessons from genetically lean mice. Annu Rev Nutr 22, 459–482. 10.1146/annurev.nutr.22.010402.102849.

30. Lim, K., Haider, A., Adams, C., Sleigh, A., and Savage, D.B. (2021). Lipodistrophy: a paradigm for understanding the consequences of “overloading” adipose tissue. Physiol Rev 101, 907–993. 10.1152/physrev.00032.2020.

31. Friedman, J.M. (2019). Leptin and the endocrine control of energy balance. Nat Metab 1, 754–764. 10.1038/s42255-019-0095-y.

32. Shimomura, I., Hammer, R.E., Ikemoto, S., Brown, M.S., and Goldstein, J.L. (1999). Leptin reverses insulin resistance and diabetes mellitus in mice with congenital lipodystrophy. Nature 401, 73–76. 10.1038/43448.

33. Savage, D.B. (2009). Mouse models of inherited lipodystrophy. Dis Model Mech 2, 554–562. 10.1242/dmm.002907.

34. Garg, A. (2004). Acquired and inherited lipodystrophies. N Engl J Med 350, 1220–1234. 10.1056/NEJMra025261.

35. Wang, M.Y., Chen, L., Clark, G.O., Lee, Y., Stevens, R.D., Ilkayeva, O.R., Wenner, B.R., Bain, J.R., Charron, M.J., Newgard, C.B., and Unger, R.H. (2010). Leptin therapy in insulin-deficient type I diabetes. Proc Natl Acad Sci U S A 107, 4813–4819. 10.1073/pnas.0909422107.

36. Maffei, M., Halaas, J., Ravussin, E., Pratley, R.E., Lee, G.H., Zhang, Y., Fei, H., Kim, S., Lallone, R., Ranganathan, S., and, et al. (1995). Leptin levels in human and rodent: measurement of plasma leptin and ob RNA in obese and weight-reduced subjects. Nat Med 1, 1155–1161. 10.1038/nm1195-1155.

37. Smith, S.J., Cases, S., Jensen, D.R., Chen, H.C., Sande, E., Tow, B., Sanan, D.A., Raber, J., Eckel, R.H., and Farese, R.V., Jr. (2000). Obesity resistance and multiple mechanisms of triglyceride synthesis in mice lacking Dgat. Nat Genet 25, 87–90. 10.1038/75651.

38. Chen, H.C., Smith, S.J., Ladha, Z., Jensen, D.R., Ferreira, L.D., Pulawa, L.K., McGuire, J.G., Pitas, R.E., Eckel, R.H., and Farese, R.V., Jr. (2002). Increased insulin and leptin sensitivity in mice lacking acyl CoA:diacylglycerol acyltransferase 1. J Clin Invest 109, 1049–1055. 10.1172/JCI14672.

39. Chen, H.C., Jensen, D.R., Myers, H.M., Eckel, R.H., and Farese, R.V., Jr. (2003). Obesity resistance and enhanced glucose metabolism in mice transplanted with white adipose tissue lacking acyl CoA:diacylglycerol acyltransferase 1. J Clin Invest 111, 1715–1722. 10.1172/JCI15859.

40. Chen, H.C., Rao, M., Sajan, M.P., Standaert, M., Kanoh, Y., Miura, A., Farese, R.V., Jr., and Farese, R.V. (2004). Role of adipocyte-derived factors in enhancing insulin signaling in skeletal muscle and white adipose tissue of mice lacking Acyl CoA:diacylglycerol acyltransferase 1. Diabetes 53, 1445–1451. 10.2337/diabetes.53.6.1445.

41. Inagaki, T., Dutchak, P., Zhao, G., Ding, X., Gautron, L., Parameswara, V., Li, Y., Goetz, R., Mohammadi, M., Esser, V., et al. (2007). Endocrine regulation of the fasting response by PPARalpha-mediated induction of fibroblast growth factor 21. Cell Metab 5, 415–425. 10.1016/j.cmet.2007.05.003.

42. Kwon, M.M., O’Dwyer, S.M., Baker, R.K., Covey, S.D., and Kieffer, T.J. (2015). FGF21-Mediated Improvements in Glucose Clearance Require Uncoupling Protein 1. Cell Rep 13, 1521–1527. 10.1016/j.celrep.2015.10.021.

43. Shih, M.Y., Kane, M.A., Zhou, P., Yen, C.L., Streeper, R.S., Napoli, J.L., and Farese, R.V., Jr. (2009). Retinol Esterification by DGAT1 Is Essential for Retinoid Homeostasis in Murine Skin. J Biol Chem 284, 4292–4299. 10.1074/jbc.M807503200.

44. Chitraju, C., Mejhert, N., Haas, J.T., Diaz-Ramirez, L.G., Grueter, C.A., Imbriglio, J.E., Pinto, S., Koliwad, S.K., Walther, T.C., and Farese, R.V., Jr. (2017). Triglyceride Synthesis by DGAT1 Protects Adipocytes from Lipid-Induced ER Stress during Lipolysis. Cell Metab 26, 407–418 e403. 10.1016/j.cmet.2017.07.012.

45. Tschop, M.H., Speakman, J.R., Arch, J.R., Auwerx, J., Bruning, J.C., Chan, L., Eckel, R.H., Farese, R.V., Jr., Galgani, J.E., Hambly, C., et al. (2011). A guide to analysis of mouse energy metabolism. Nat Methods 9, 57–63. 10.1038/nmeth.1806.

46. Folch, J., Lees, M., and Sloane Stanley, G.H. (1957). A simple method for the isolation and purification of total lipides from animal tissues. J Biol Chem 226, 497–509.

47. Taguchi, R., and Ishikawa, M. (2010). Precise and global identification of phospholipid molecular species by an Orbitrap mass spectrometer and automated search engine Lipid Search. J Chromatogr A 1217, 4229–4239. 10.1016/j.chroma.2010.04.034.

